# Ancestral complexity and constrained diversification of the ant olfactory system

**DOI:** 10.1101/2024.10.03.616251

**Authors:** Simon Marty, Antoine Couto, Erika H. Dawson, Neven Brard, Patrizia d’Ettorre, Stephen H. Montgomery, Jean-Christophe Sandoz

**Affiliations:** Evolution Genomes Behavior and Ecology (UMR9191), IDEEV, Université Paris-Saclay, CNRS, IRD, Gif-sur-Yvette 91190, France; School of Biological Sciences, University of Bristol, Bristol Life Sciences Building, 24 Tyndall Avenue, Bristol, BS8 1TQ, UK; Laboratory of Experimental and Comparative Ethology, UR 4443 (LEEC), Université Sorbonne Paris Nord, 99 avenue J.-B. Clément, Villetaneuse 93430, France; Institut Universitaire de France (IUF), 103 Boulevard Saint-Michel, Paris 75005, France

**Author notes:** S.M. and A.C. contributed equally to this work. S.H. M. and J.-C. S. contributed equally to this work.

## Abstract

Ants are a monophyletic but diverse group of social insects whose heightened olfactory ability has been crucial to their evolutionary success. Their complex olfactory system is believed to have evolved due to the expansion of a specialized olfactory subsystem and the associated clade of olfactory receptors. Specifically, ants exhibit specialized antennal hairs known as basiconic sensilla, whose neurons project to a distinctive cluster of numerous, small glomeruli in their antennal lobes. This adaptation is believed to be linked to their social lifestyle, enabling the detection of recognition cues like cuticular hydrocarbons (CHCs), which are essential for nestmate recognition and maintaining colony cohesion. However, our understanding of the ant olfactory system remains incomplete, lacking evolutionary context and phylogenetic breadth, which leaves the complexity in their most recent common ancestor uncertain. We thus conducted a comparative study of neuroanatomical traits across the phylogeny of the Formicidae. Our findings reveal a common blueprint for the ant olfactory pathway, alongside lineage-specific adaptations. This highlights a dynamic evolution, particularly for the CHC-related subsystem. Ancestral trait reconstructions indicate that olfactory sophistication predates the most recent common ancestor of ants. Additionally, we found that the chemical complexity of species-specific recognition cues is associated with neuronal investment within the olfactory system. Lastly, behavioral experiments on anatomically divergent ant species show that, despite variation in neuroanatomical traits, ants consistently discriminate nestmates from non-nestmates. This suggests that the evolution of ants’ olfactory system integrates sensory adaptations to diverse chemical environments, facilitating communication, a key to social behaviors.

## Introduction

Communication is a cornerstone of social living. Through the exchange of information, group members resolve conflicts, align goals, and synchronize efforts, ultimately facilitating cooperation. Consequently, whether in mammalian societies or vast colonies of social insects, social evolution is believed to influence the development of recognition and communication systems, thereby supporting reliable social interactions (1-4). Despite significant progress in understanding the neural components of communication, particularly in social insects (5), a notable knowledge gap remains regarding the evolution of the sensory systems that support this critical function. Therefore, exploring the roles of novel neuronal populations and other adaptations within the sensory pathways that support communication, is key to understanding the mechanisms underlying the evolution of social behavior.

Ants stand as prominent models of group living and cooperation, owing to their remarkably large and complex colony organizations, diverse kin structures, and extensive interspecific variation in morphological traits, dietary preferences, foraging behaviors, and life history strategies (6, 7). At the core of their cooperative behaviors lies a sophisticated communication system that supports pheromonal signaling and the perception of recognition cues, primarily odorant compounds detected by the olfactory system (8, 9). Accordingly, ants possess one of the most complex olfactory systems among insects (10-12), as evidenced by studies on selected species which highlighted its pivotal role in social interactions and scent-guided behaviors (13, 14). Nevertheless, a significant gap remains regarding the evolutionary trajectory of ants’ olfactory system. Understanding these trajectories could illuminate the adaptive significance of olfaction within the context of sociality and the remarkable taxonomic radiation of ants.

Insects’ antennae are typically covered with different types of sensory hairs known as sensilla, which enclose the dendrites of olfactory sensory neurons (OSNs). The axons of these neurons project to the antennal lobe (AL) in the brain, where they form glomeruli, discrete spherical structures that serve as processing units. Each OSN generally expresses a single olfactory receptor (OR) – together with the ubiquitous co-receptor – which defines its response profile to odorant stimuli (15, 16). The OR expression also dictates the specific glomerular target of each neuron, establishing a nearly one-to-one correspondence between ORs and glomeruli (17). These olfactory glomeruli serve as central hubs where local interneurons and neuromodulatory neurons refine olfactory information before it is relayed to higher brain centers by projection neurons (18).

The complexity of ants’ olfactory system is supported by two key observations. First, ants possess a high number of AL glomeruli and a greater abundance of associated OR genes compared to most other insect clades (10, 11, 15, 19). Second, ants exhibit an olfactory specialization which has been associated with the detection of recognition cues (5). This specialized pathway is characterized by a unique type of antennal sensilla, the basiconic sensilla, wherein OSNs express a specific clade of 9-exon ORs (11) and project exclusively to a distinct cluster of glomeruli (20, 21), which lack serotoninergic innervation (22-25). Notably, electrophysiological recordings have revealed that cuticular hydrocarbons (CHCs), which are identity-signaling compounds, are sensed by the 9-exon ORs (26-28) and basiconic sensilla OSNs (29-33). These molecules are pivotal in insect communication (34), conveying information about species, colony affiliation, and reproductive status (35), thereby facilitating the recognition of nestmates over non-nestmates and the maintenance of colony cohesion (36). Thus, this olfactory subsystem is believed to have evolved and expanded significantly in ants, driven by the need to meet sophisticated communication demands within complex social colonies (11, 19, 28, 37).

At this stage, our knowledge of ants’ olfactory system is biased towards a few specific clades, significantly limiting insight into how its sophistication relates to ants’ social behavior and ecology. For example, it is not yet known whether the increased complexity of the AL and the extensive repertoire of ORs translate into enhanced accuracy in recognizing social identities, such as distinguishing between nestmates and non-nestmates. Additionally, the diversification of recognition cues, particularly the chemical composition and complexity of CHC profiles (38, 39), likely interplays with the evolution of the ant olfactory subsystem. This co-evolution suggests a dynamic feedback mechanism in which increasingly complex chemical signals drive adaptations within the neural and sensory apparatus. However, to date, little is known about the influence of dynamic communication demands and varied social structures on the evolutionary trajectory of the olfactory system across the ant phylogeny (40). Therefore, it remains uncertain whether the olfactory system was already complex at the onset of ants’ remarkable taxonomic radiation or if its complexity evolved during their diversification.

Here, we employed a comparative approach to explore the evolutionary history of key olfactory neuroanatomical traits in ants, examining whether variation in ecology and social structure have influenced differential investment in their olfactory system. Overall, we investigated 14 species from 8 Formicidae subfamilies, differing in dietary behavior, ecological niche, and colony kin structure. We compared the distribution of basiconic sensilla across antennal segments and, within the antennal lobe, examined neuromodulatory populations to identify the glomeruli of the ant’s olfactory subsystem involved in social recognition. Additionally, we analyzed neuropil volumes and glomerular counts, integrating these measurements to provide a comprehensive assessment of the olfactory organization in ants. We conducted ancestral state reconstructions and evolutionary rate analyses on the number of glomeruli to identify derived traits and periods of rapid evolution. Using this evolutionary framework, we finally tested whether the anatomical variation observed in the olfactory system correlates with behavioral differences between species, and are linked to ecological or chemical factors. By integrating these diverse approaches, we aim to elucidate the role of ecological pressures and diverse social structures in shaping the evolution of the olfactory system across the ant phylogeny.

## Results

### Diversity and distribution of antennal sensilla

To explore the evolution of sensory structures in ants, we conducted scanning electron microscopy on the antennae of 13 ant species spanning a broad phylogenetic range. Among the various morphological types of sensilla, basiconic sensilla, identified by their peg-in-socket shape and porous tip, contrast with other sensory hairs (41). This sensillum type was consistently present across all studied species, exhibiting only slight morphological differences (Fig. 1A-C).

**Figure 1.**
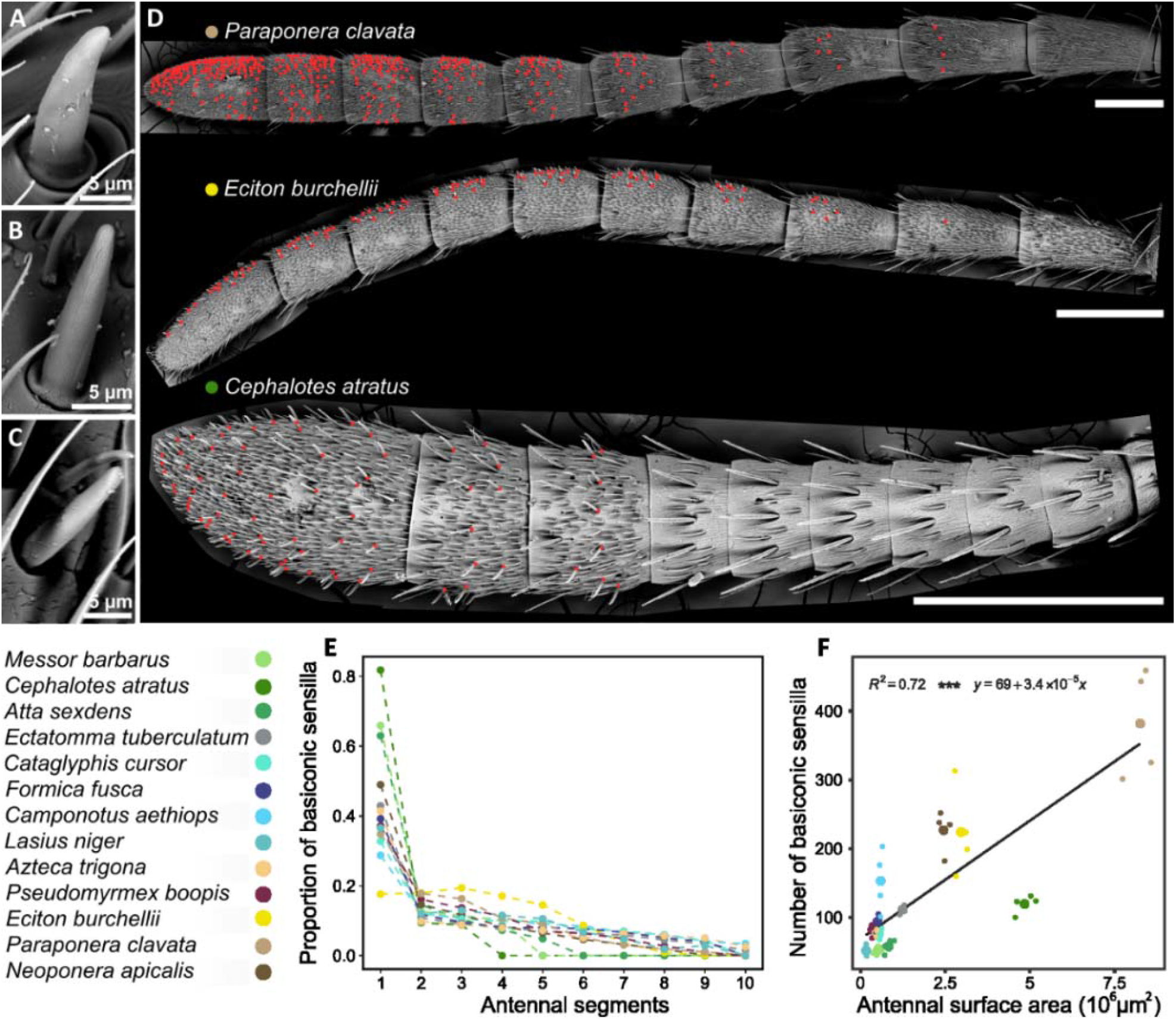
Distribution of basiconic sensilla on ant antennae. (A-C) Scanning electron micrographs of basiconic sensilla across ant species. These sensilla show slight morphological differences, including a reduction in base thickness from basal taxa (*Neoponera apicalis*, A) to Formicinae (*Formica fusca*, B) and Myrmicinae clades (*Cataglyphis cursor*, C). (D) Scanning electron micrographs of *Paraponera clavata, Eciton burchellii*, and *Cephalotes atratus* antennae, illustrating the distribution of basiconic sensilla (red dots) across antennal segments. These sensilla are more densely packed on the distal segments of the antenna. Scale bars are set at 0.5 mm. (E) Relative proportion of basiconic sensilla across antennal segments in various ant species (colored dots). Basiconic sensilla are notably concentrated in the distal part of the antennae, prominently in Myrmicinae (varying green shades). (F) Number of basiconic sensilla on the antenna plotted against antenna surface area, showing species means (large colored dots) and individual data points (small dots). The number of basiconic sensilla correlates strongly with the size of the antenna (Pearson test: t = 4.97, df = 11, p < 0.001, R^2^ = 0.72).

In ants, the distribution of basiconic sensilla is strongly biased towards the distalmost segments of the antennae (Fig. 1D-E, two-way ANOVA, p < 0.001 across flagellomeres), with their proportion decreasing towards proximal segments. Consequently, there is a progressive decline in both the number and density of sensilla along the antenna (Fig. S1 D, E), albeit with significant variability in attenuation patterns across species (Fig. 1E, two-way ANOVA, *species* x *segment* interaction, p < 0.001, Fig. S1D, E). The decline is particularly pronounced in Myrmicinae species, where basiconic sensilla are absent beyond the third segment in *Cephalotes atratus*, the fourth segment in *Atta sexdens*, and the fifth segment in *Messor barbarus*. In contrast, *Eciton burchellii* (Dorylinae), exhibits a more even distribution of basiconic sensilla across the first four segments, with each containing approximately 18% of the total count (Fig. 1D, E, post-hoc Tukey, p > 0.05 between segments 1, 2, 3, and 4). Across species, the total number of basiconic sensilla varies significantly (Table S1 and Fig. S1E, two-way ANOVA, p < 0.001), ranging from 52.2 ± 5.6 in *M. barbarus* to 382 ± 80 in *Paraponera clavata*. These differences primarily reflect variation in antenna size, as indicated by the strong correlation between basiconic sensilla counts and the measured antennal surface area (Fig. 1F; Pearson test, t = 4.9664, df = 11, p < 0.001, R^2^ = 0.72).

### Characterization of anatomical regionalization in the antennal lobe

Given the marked variation in sensilla numbers, despite a consistent organizational pattern, we investigated how these differences affect the structure of the antennal lobe (AL). Using immunohistochemistry and confocal microscopy, we characterized the AL across 14 ant species.

In all examined species, the AL exhibits two distinct regions resembling glomerular rings (Fig. 2A-C). The dorso-caudal region, featuring more compact glomeruli compared to the rostral region of the AL (referred to as the Main-AL, Fig. 2A-C), corresponds to the T_B_ cluster (20, 21). Basiconic sensilla OSNs project exclusively into this cluster (20, 21), which is reported to lack serotonin innervation in the *Camponotus* clade, unlike the rest of the AL typically innervated by serotonergic neurons (22, 23). Lack of serotonergic innervation has been proposed as a marker for the T_B_ cluster (42). Therefore, we investigated the innervation patterns of serotonergic neurons across the Formicidae, as a marker of functional and anatomical regionalization within the AL of ants. We found that the T_B_ cluster lacks serotonergic neurons in *A. sexdens* (Fig. S2A), *M. barbarus* (Fig. 2A), *C. atratus, Aphaenogaster senilis*, and *Camponotus aethiops* (Fig. S2B), while the rest of their ALs exhibit clear innervation (summarized in Fig. 2D). Similarly, the T_B_ cluster in *P. clavata* and *E. burchellii* appears to lack serotonergic fibers, although this observation is less certain due to lower quality of our immunostaining replicates. In contrast, *Ectatomma tuberculatum* (Fig. 2B), *Neoponera apicalis* (Fig. 2SC), *Pseudomyrmex boopis* (Fig. 2SD), *Formica fusca, Lasius niger, Cataglyphis cursor*, and *Azteca trigona* all exhibit clear serotonergic innervation in the T_B_ cluster (as summarized in Fig. 2D), with varying densities across both the Main-AL and the T_B_ glomeruli.

**Figure 2.**
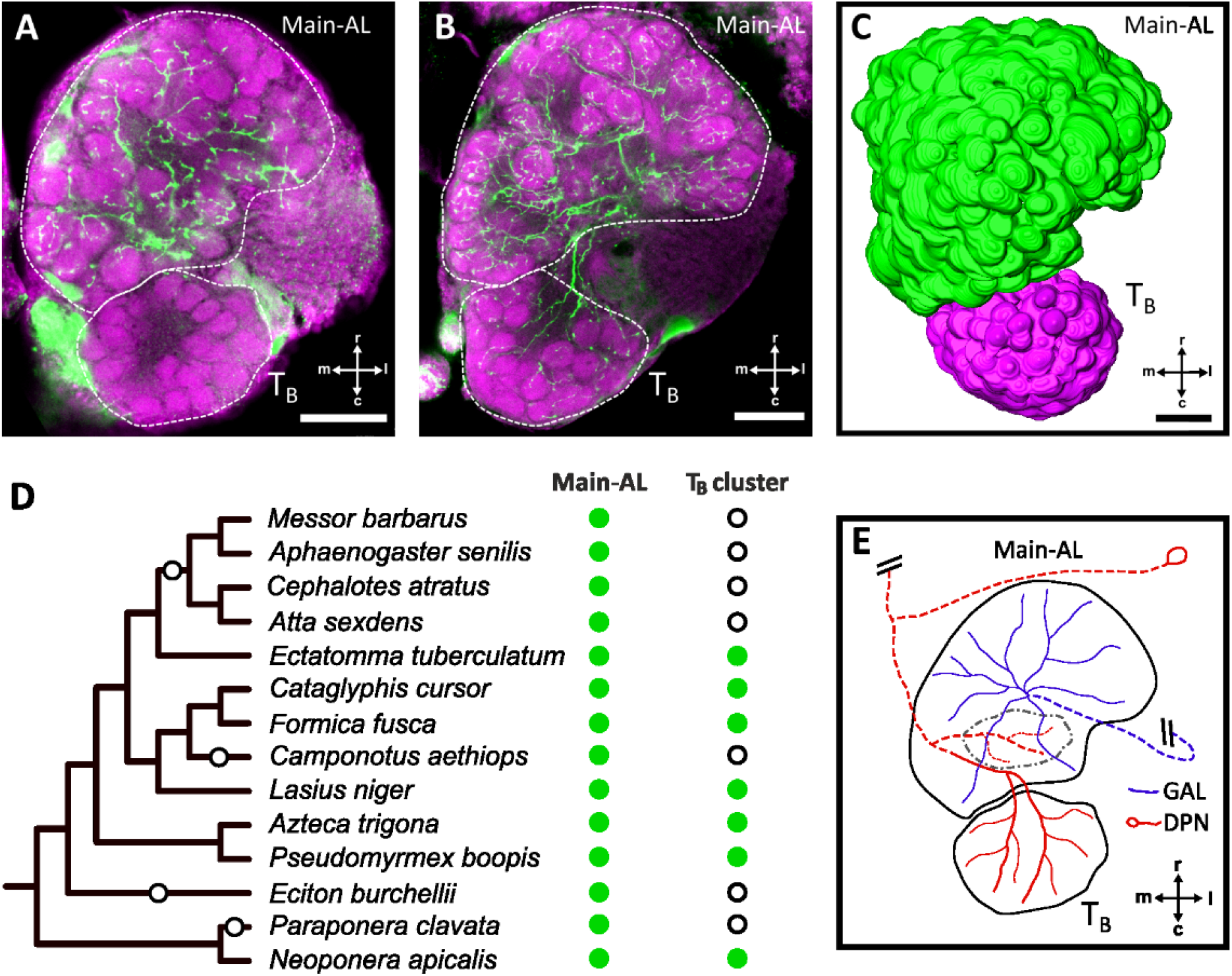
Anatomical characterization of the antennal lobe. (A, B) Confocal optical sections of the antennal lobe in A) an ant species lacking serotonin innervation in the T_B_ cluster (*Messor barbarus*) and B) a species with serotonin immunoreactivity in this region (e.g., *Eciton tuberculatum*). The glomeruli are stained with hydrazide conjugated dye displayed in magenta, and immunolabeled serotonergic projections are displayed in green. The AL subregions are outlined with dashed lines. (C) 3D reconstruction of glomerular volumes and antennal lobe regionalization in *Neoponera apicalis*. The Main-AL, which receives serotonergic innervation, is shown in green, while the T_B_ cluster, which lacks such innervation in some species, is in magenta. (D) Variation in serotonergic innervation across ant species summarized on the phylogeny. Green dots indicate serotonergic innervation, while empty circles in the T_B_ cluster column represent the absence of such projections. Losses of serotonergic projections, predicted by the most parsimonious scenario, are marked with empty circles on the tree branches. (E) Diagram summarizing serotonergic projections in the AL of ants with serotonergic T_B_ clusters. In all species, the giant neuron innervating the AL (GAL, in blue) innervates the Main-AL, while the deutocerebral projection neuron (DPN, in red) targets the dorsal cluster of glomeruli (black dashed lines). In species with immunoreactive T_B_ clusters, DPN extends an additional branch to innervate the T_B_ glomeruli. Dashed lines represent structures that are dorsal relative to the AL. All scale bars represent 50 μm (r, rostral; c, caudal; m, medial; l, lateral).

We further traced and examined the innervation pattern of serotonin-immunoreactive neurites within the two subregions of the AL (Fig. 2E). In all species, the glomeruli of the Main-AL primarily receive projections from a neuron known as the giant neuron (GAL), which originates in the subesophageal zone and innervates the AL (23). Additionally, the soma of another neuron, known as the deutocerebral projection neuron (DPN), is consistently observed within the lateral cell cluster near the rostral edge of the AL (23). In all species, this DPN innervates a few glomeruli in the dorsal region of the Main-AL and sends ipsilateral projections towards the mushroom bodies (Fig. 2E). However, in T_B_ immunoreactive species, we observed that the DPN exhibits an additional branch that specifically innervates the T_B_ cluster. Therefore, the absence of serotonergic innervation in T_B_ clusters correlates with the loss of this DPN extension, which can serve, along with morphological characteristics, as a key anatomical feature for characterizing T_B_ across species.

### Volumetric relationships in the ant antennal lobes

Using 3D models reconstructed from confocal image stacks (Fig 2C), we measured the volume of the AL as the total glomerular area and found strong differences across species (Table S1, Fig. S2E, Kruskal-Wallis test, χ^2^ = 38.9, df = 14, p < 0.001). This is illustrated by the 20-fold difference in AL size between *L. niger* (0.69 × 106 µm3) and *P. clavata* (12.1 × 106 µm3), likely reflecting their substantial difference in body size. We also investigated neural investment in the two AL subdivisions, which are believed to serve different functions, by examining the scaling relationship between the volume of the Main-AL and that of the T_B_ cluster. Despite considerable total volumetric variation, we found a significant correlation between the volumes of the Main-AL and that of the T_B_ cluster (Fig. 3A; Pearson test, t = 15.8, p < 0.001, R^2^ = 0.86). This indicates that neuronal investment in the T_B_ cluster follows a relatively stable allometric relationship across ant species (Table S1).

**Figure 3.**
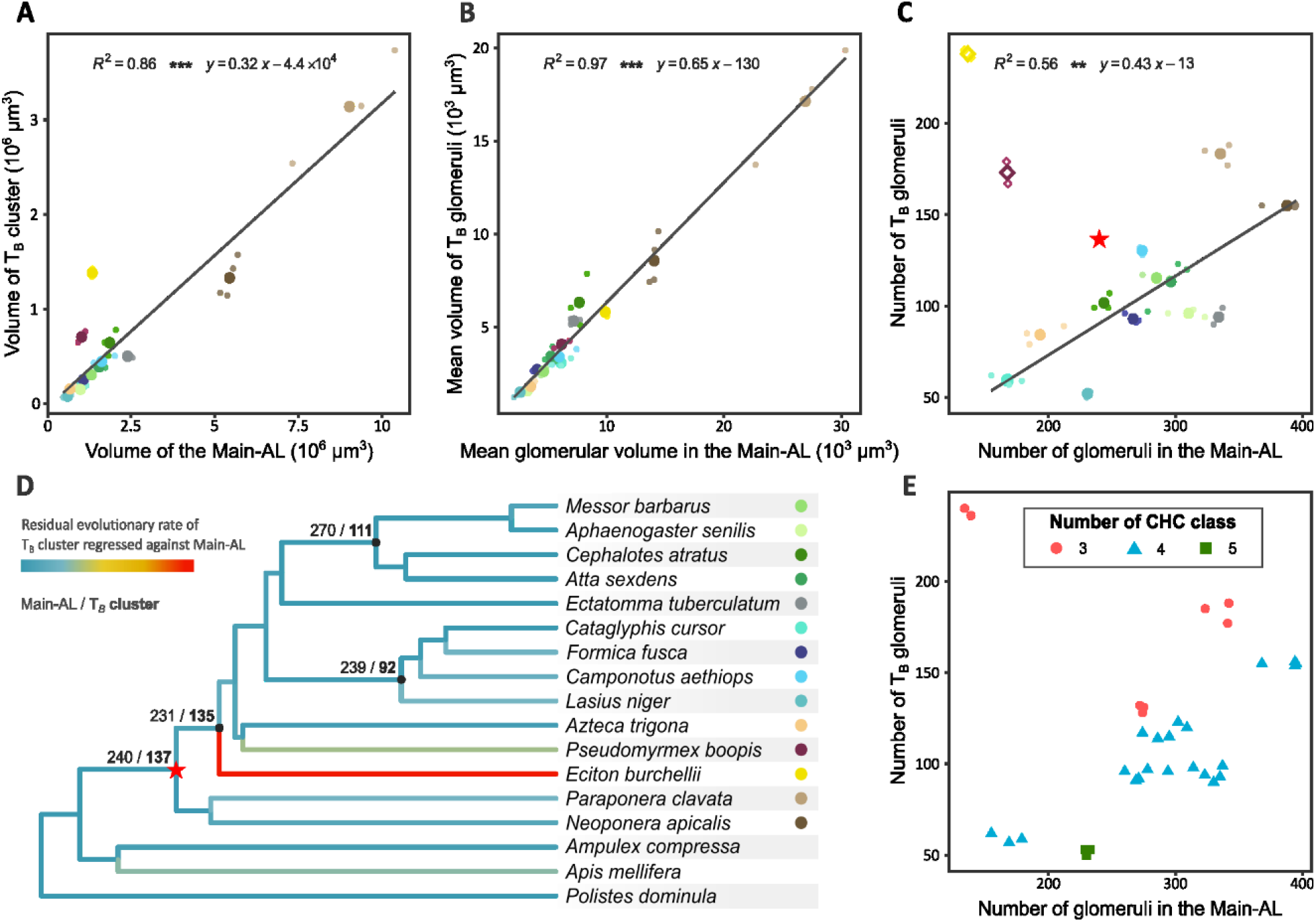
Evolutionary dynamics in the ant antennal lobe. (A) Volume of the T_B_ cluster plotted against the volume of the Main-AL across different ant species. There is a significant correlation between the volumes of the T_B_ cluster and of the Main-AL (t = 15.8, p < 0.001). (B) Mean volume of glomeruli in the T_B_ cluster plotted against the mean volume of glomeruli in the Main-AL across different species. There is a correlation between the mean glomerular volumes in the two subregions of the AL (t = 33.9, p < 0.001), with T_B_ glomeruli being consistently smaller than those of the Main-AL. (C) Number of glomeruli in the T_B_ cluster plotted against the number of glomeruli in the Main-AL. These counts are not correlated (t = 0.70, p = 0.48). However, when excluding outlier species (squares), the counts significantly correlate (t = 6.61, p < 0.01). The red star on the graph represents the putative position of ant’s most recent common ancestor (see text and Fig S3A-C). (D) Residual evolutionary rates of the number of glomeruli in the T_B_ cluster, relative to the Main-AL, mapped onto the branches of the phylogenetic tree of the sampled species. The branches leading to *Eciton burchellii* and, to a lesser extent, *Pseudomyrmex boopis*, display high evolutionary rates, suggesting two independent expansions of the T_B_ cluster. Numbers indicated at the nodes represent the inferred ancestral state estimates for the number of glomeruli in both the Main-AL and the T_B_ cluster. (E) Plot showing the number of glomeruli, similar to (C), with species categorized according to the number of compound classes (alkanes, alkenes, mono-, di- or tri-methyl alkanes) within their CHC profiles. There is a significant effect of the number of CHC classes on the proportion of glomeruli between the T_B_ cluster and the Main-AL (*pMCMC* < 0.01).

We examined the relationship between the mean glomerular size within both subregions, by dividing each subregion’s volume by its number of glomeruli. Our analysis revealed a significant correlation between the volumes of glomeruli in the T_B_ cluster and those in the Main-AL (Fig. 3B; Pearson test, t = 33.9, p < 0.001, R^2^ = 0.97). The slope (0.65, [0.607; 0.684] with 95% CI) is significantly lower than 1 (t test, t = -32.3, p < 0.001), with a negative y-intercept. This indicates that, across species, glomeruli of the T_B_ cluster are consistently smaller than those of the Main-AL.

### Glomerular counts and evolutionary dynamics in the ant antennal lobes

The glomerular count offers a measure of the AL’s computational power, with more glomeruli suggesting a greater capacity to integrate and discriminate diverse olfactory information through combinatorial processing (43). Across our sample set, the number of glomeruli in the AL of worker ants varied significantly across species (Table S1 and Fig. S3A; Kruskal-Wallis test, χ^2^ = 38.5, df = 14, p < 0.001), ranging from 227 ± 10 (mean ± SD) glomeruli in *C. cursor* to 543 ± 13 glomeruli in *N. apicalis*. These differences are evident within each AL subdivision, with significant variation in the number of glomeruli in both the T_B_ cluster (Table S1 and Fig. S3B, C; Kruskal-Wallis test, χ^2^ = 38.9, df = 14, p < 0.001) and the Main-AL (Table S1 and Fig. S3B, C; Kruskal-Wallis test, χ^2^ = 39.4, df = 14, p < 0.001).

We further examined whether ants invest differently in the two AL subdivisions by studying the scaling relationship between the number of glomeruli in the T_B_ cluster and the Main-AL. These counts are not correlated across all ants (Pearson test, t = 0.70, p = 0.48), due to notable deviations of a few species (Fig. 3C). We therefore analyzed the evolutionary rate of glomerular numbers using a variable-rate Brownian motion model across the ant phylogeny (44). The tree was rooted using additional glomerular counts from the paper wasp *Polistes dominula* (Fig. S2F), the emerald cockroach wasp *Ampulex compressa* (Fig. S2G), and the honeybee *Apis mellifera* ((45), see Suppl methods). We generated separate models for the two subregions, and displayed the T_B_ cluster residuals regressed against the Main-AL along the tree to identify deviations in the allometric relationship between these subregions (Fig. 3D). Our analysis revealed a higher evolutionary rate on the branches leading to *E. burchellii* and, to a lesser extent, *P. boopis*, compared to the rest of the tree. These two species are notable outliers, with their T_B_ clusters comprising 63.3% and 50.8% of the total AL glomeruli, respectively. Excluding these species, the glomeruli numbers between the two AL subdivisions show a significant correlation (Fig. 3C; Pearson test, t = 6.61, p < 0.01, R^2^ = 0.56). This suggests that the variation in glomerular numbers is allometrically consistent between AL compartments, except in *E. burchellii* and *P. boopis*, which exhibit a disproportionately high number of T_B_ glomeruli. Incorporating literature data on *Ooceraea biroi* (11) and *Dolichoderus sp*. (46), we corroborate this trend and provide support for an expanded T_B_ cluster in the Dorylinae subfamily (Fig. S3D, E).

We finally used maximum likelihood ancestral state estimation to reconstruct the evolutionary history of the AL across ants. The global model predicted approximately 367 AL glomeruli in the most recent common ancestor (MRCA) of ants (Fig. S3A). Specifically, the analysis predicted 137 glomeruli for the T_B_ cluster (Fig. 3D, and Fig. S3B) and 242 for the Main-AL (Fig. 3D, and Fig. S3C) in independent models. Contrary to previous conclusions (11, 37), our findings suggests that the T_B_ subsystem was already elaborated before the extensive taxonomic radiation of ants. Furthermore, the model estimated 142 T_B_ glomeruli in the MRCA of Dorylinae and other formicoids, and 133 T_B_ glomeruli in the MRCA of *P. boopis* and the Dolichoderinae, indicating distinct expansions of T_B_ glomeruli relative to the Main-AL in these lineages (Fig 3D and Fig. S3B, D). These expansions were accompanied by a reduction in the number of other glomeruli (Fig. S3C), distinguishing these species from the general trend.

### Social and chemical predictors of glomerular variation

We next aimed to elucidate the social and ecological correlates of the number of glomeruli in the T_B_ cluster and the Main-AL using phylogenetically controlled mixed models, which incorporate social and chemical predictors (Table S2). First, we analyzed social predictors, including types of polygyny (strictly monogynous versus facultatively or obligatorily polygynous), foraging strategies (solitary versus collective), and colony sizes (categorized as less than 1000, between 1000 and 10000, and more than 10000). None of these factors had any significant effect on the number of glomeruli (total, T_B_ or Main AL), the number of basiconic sensilla, or volumetric measures of the AL (total, T_B_ or Main AL) (see SuppI-Data file). Likewise, these social factors did not affect the scaling of glomeruli between the T_B_ cluster and the Main-AL (Fig. S4; polygyny: *p*_*MCMC*_ = 0.388; foraging: *p*_*MCMC*_ = 0.827; colony size: *p*_*MCMC*_ = 0.386 and 0.387).

Next, we investigated whether variation in species-specific cuticular chemical profiles is related to these anatomical traits, using chemical analyses by gas-chromatography coupled with mass-spectrometry (see Supplementary Material, 1.7 Chemical analysis of CHC profiles). From each species’ chemical profile, we extracted three key variables: the number of individual CHCs (CHC number), the number of distinct CHC classes (ranging from 1 to 5, including alkanes, alkenes, mono-, di-, or tri-methyl alkanes), and a measure of profile complexity, the Shannon Index (SI; see methods).

All models that included the SI showed it as a significant predictor of the number of glomeruli in the T_B_ cluster (full model: posterior mean = -2.014, *p*_*MCMC*_ = 0.005). Specifically, CHC profile complexity was negatively associated with the number of T_B_ glomeruli, suggesting that species with less complex CHC profiles have more glomeruli in the T_B_ cluster. However, the inclusion of the SI only marginally improved the model fit compared to the model in which it was excluded (ΔDIC = 0.25).

Since the Shannon Index accounts for both the number of CHC compounds and the number of classes, we explicitly tested these parameters in the model. While the number of compounds did not significantly influence the number of T_B_ glomeruli (posterior mean = 0.0049, *p*_*MCMC*_ = 0.595), the number of CHC classes was negatively associated with this count (posterior mean = -0.827, *p*_*MCMC*_ = 0.045). This indicates that a lower diversity of CHC classes is associated with a higher number of T_B_ glomeruli, although the improvement in DIC was minimal when comparing the full model to the one without CHC class (ΔDIC = 0.05). However, none of the chemical variables (SI, CHC number, CHC classes) were significant predictors of glomeruli numbers in the Main-AL or of the total glomerular count (for SI, Main: posterior mean = -0.373, p_*MCMC*_ = 0.591; total: posterior mean = -0.873, p_*MCMC*_ = 0.0976). These chemical factors also did not significantly predict the volume of the AL (total, T_B_, or Main AL) or the number and density of basiconic sensilla. Investigating the scaling of glomeruli numbers between the T_B_ cluster and the Main-AL, we again found SI to be significantly associated with investment in both olfactory regions (posterior mean = -1.924, *p*_*MCMC*_ = 0.0100).

In conclusion, models including chemical predictors were the most parsimonious and best-fitting, though they showed only a slight DIC improvement (ΔDIC < 1). Specifically, the chemical variables, SI and the number of CHC classes, emerged as significant predictors of ants’ investment in glomeruli numbers in the T_B_ cluster.

### Neuroanatomy’s impact on ants’ discrimination performance

Building on the interspecific differences we found in serotonergic innervation and glomerular number, we tested whether this variation reflects differences at the behavioral level. We conducted a nestmate discrimination assay on a selected set of species: *M. barbarus* and *F. fusca*, which share approximately the same number of glomeruli in both subregions, and *L. niger*, which has a lower number of glomeruli in the T_B_ cluster compared to the other two species. Additionally, the T_B_ clusters of *F. fusca* and *L. niger* receive serotonergic innervation, while it is absent in *M. barbarus*. Workers of all these three species were able to perform a discrimination task, consistently showing higher aggression levels towards non-nestmates than towards nestmates (Fig. 4A; *F. fusca*: χ^2^ = 18.2, df = 1, p < 0.001; *L. niger*: χ^2^ = 5.52, df=1, p = 0.0188; *M. barbarus*: χ^2^ = 13.0, df = 1, p < 0.001). Overall, we found that the three species were similarly efficient at discriminating between nestmate and non-nestmate stimuli, with differentiation scores that did not significantly differ among species (Fig. 4B; χ^2^ = 0.098, df = 2, p = 0.95).

**Figure 4.**
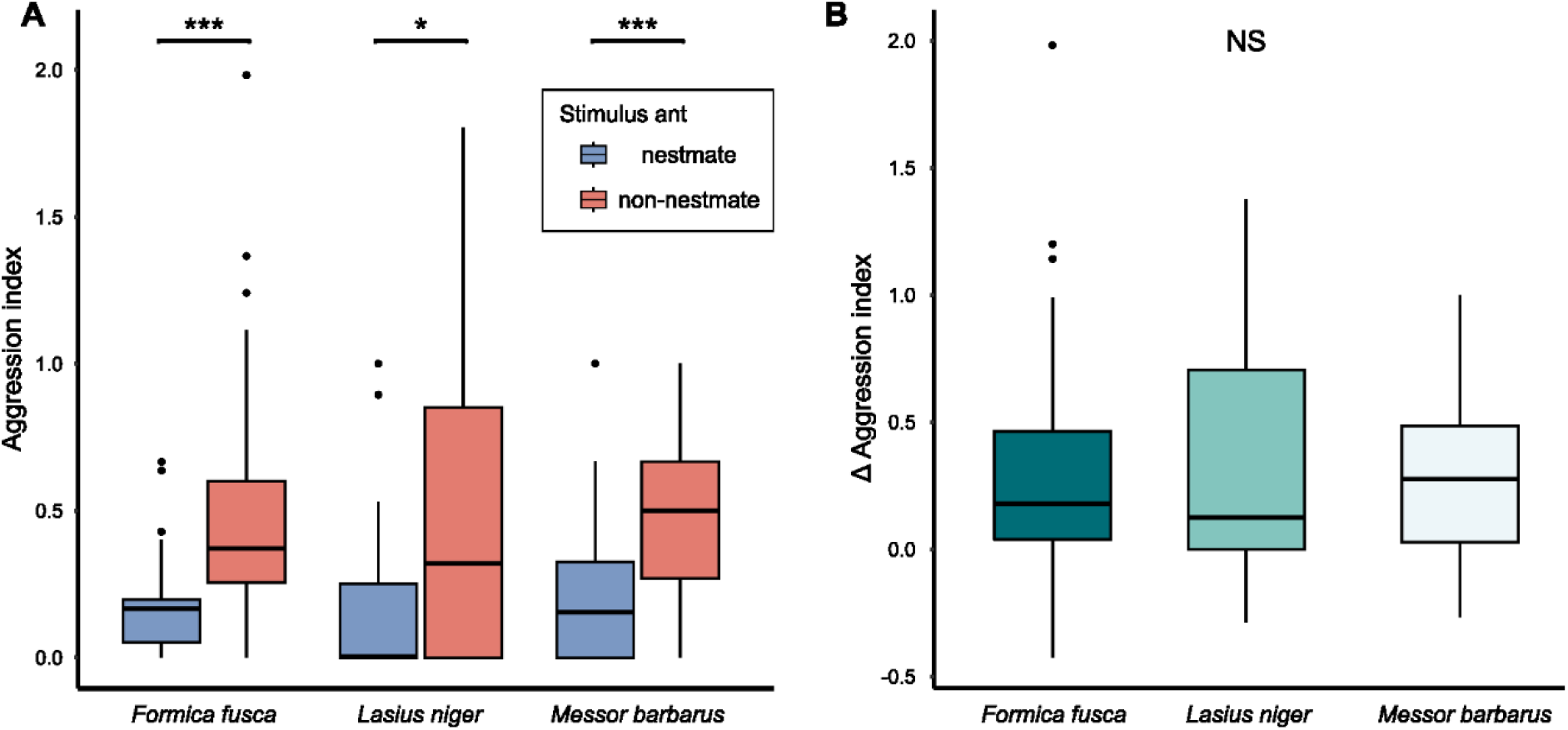
Comparative analysis of nestmate discrimination abilities. (A) Boxplots show median, quartiles, and minimum/maximum values (whiskers) for aggression index. All investigated species discriminate nestmate from non-nestmate odors (^***^: p < 0.001, ^*^: p < 0.01). (B) Boxplots represent median, quartiles, and minimum/maximum values (whiskers) for Delta aggression scores. Delta scores were calculated by subtracting the aggression index of nestmates from non-nestmates. Positive scores indicate more aggression towards non-nestmates, while negative scores indicate more aggression towards nestmates. The species were equally efficient at discriminating between nestmate and non-nestmate stimuli (NS: p > 0.05). In each boxplot, black circles indicate outliers. N = 30 individual ants for each species.

## Discussion

In this study, we investigated anatomical variation within the olfactory system in a subset of ant species, encompassing all major taxonomic lineages of the Formicidae, to explore the evolution and adaptation of their CHC-related subsystem. We observed significant variation in olfactory traits, including the number of basiconic sensilla, as well as the volume and number of glomeruli in the antennal lobes (AL). Despite this variation, the olfactory systems of these ant species exhibit a common ground plan characterized by a higher proportion of basiconic sensilla on the distalmost segments of the antennae, and consistent proportions of neuropil volume and glomeruli number between AL subregions. However, we also identified remarkable outliers, demonstrating a high evolutionary rate of T_B_ glomeruli within specific subfamilies, resulting in the expansion of the T_B_ cluster relative to the Main-AL. Nonetheless, contrary to previous assumptions (11, 19, 37), ancestral traits reconstruction suggests that the olfactory system was already sophisticated in the most recent common ancestor (MRCA) of ants, with possibly 367 glomeruli, including 142 within the CHC-related subsystem. In considering ecological and chemical factors that might influence this evolution, the complexity of ants’ CHC profiles, particularly CHC class diversity, emerged as a potential predictor of variation in glomerular investment, albeit with modest explanatory power. Nonetheless, neither the relative nor absolute glomerular investment appears to influence nestmate discrimination performance across species. Thus, our results suggest that ants possessed a sophisticated olfactory system prior to their taxonomic radiation, and subsequent evolution was likely shaped by the necessity to maintain high olfactory performance across diverse ecological conditions and life history strategies.

### An olfactory subsystem for hydrocarbon sensing in ants

Ants universally bear a wide diversity of CHCs on their body surface, which serve as efficient recognition cues (47). These compounds play a pivotal role in social communication, especially in nestmate recognition, which is crucial for preventing exploitation by competitors and parasites, and maintaining colony cohesion. Given that this task is typically carried out by female workers, the absence of basiconic sensilla and T_B_ glomeruli in males hints at their specialized function in detecting recognition cues (25, 48-50). This idea is further supported by electrophysiological studies demonstrating the role of basiconic sensilla OSNs (29, 31, 33), as well as expression-biased ORs of the 9-exon clade (11, 51), in responding to CHCs (26, 27). While this subsystem has been described in a handful of species spanning only three Formicidae subfamilies (*Camponotus spp*. (24, 50), *A. vollenweideri* (20), *O. biroi* (11), *E. burchellii* (12)), the present study provides an evolutionary perspective across the ant phylogeny, showing the presence of this subsystem in all eight major Formicidae clades.

First, we demonstrated the presence of basiconic sensilla on the antennae of all species. We found a higher density of basiconic sensilla on the distal segments of the antennae, despite varying quantities across species. Consistent with previous observations (11, 49, 52, 53), this finding supports the role of basiconic sensilla as close-range chemoreceptors, as the distal segments of the antennae come into close proximity to other individuals during antennation (54).

Remarkably, all species also exhibited a distinct separation between the Main-AL and the T_B_ cluster, forming two glomerular rings on optical sections, with the T_B_ cluster characterized by a congregation of small, uniformly sized glomeruli. The relative size of glomeruli typically reflects the quantity of incoming OSNs, which in turn correlates with detection sensitivity to associated odorants (55-57). As such, the smaller size of T_B_ glomeruli, indicating fewer OSNs inputs, likely reflects reduced sensitivity compared to the glomeruli of the Main-AL. Consequently, T_B_ glomeruli might be well adapted for close-range detection but less effective in detecting more dispersed environmental odors. Selection pressures on the specific ability to perceive CHC profiles may therefore shape the number of T_B_ glomeruli rather than their size, emphasizing discrimination over sensitivity.

### Evolutionary dynamics of ant olfactory systems

The number of ORs genes and the corresponding glomerular count in the AL are believed to reflect a species’ olfactory discrimination power, as odors are processed through the combinatorial activity of OSN populations (43). Ants demonstrate extensive variation in glomeruli numbers, from 198 in *Cataglyphis fortis* (58) to 630 in *Apterostigma cf. mayri* (10), which is at least 3-4 times higher than typical Holometabolous insects such as *Drosophila melanogaster* with 52 (59), *Aedes aegypti* with 50 (60), or *Manduca sexta* with 65 (61). This high number is believed to heighten ants’ olfactory discrimination abilities, supporting their prominent collective behaviors (40). Nonetheless, such substantial glomerular counts might have either expanded in response to increasing social complexity or, conversely, been a pre-existing feature that facilitated the radiation of social lineages (2, 62). Our ancestral state reconstructions of glomerular number suggest that an estimated 383 glomeruli were already present in the MRCA of ants, supporting the latter hypothesis. From this ancestral state, our analyses suggest that the number of glomeruli evolved dynamically to adapt to the ecological conditions of different clades, particularly the variety of chemical cues present in their environment, rather than social factors such as polygyny, colony size, foraging strategy.

The specific family of 9-exon ORs has been highlighted as particularly expanded in ants, and showing signatures of positive selection (11, 19, 37), suggesting that dynamic gene family evolution has accompanied the evolution of ant sociality. Additionally, several pieces of evidence hint that the size of the 9-exon OR repertoire and the associated number of glomeruli within the T_B_ cluster may adapt to varying social traits. For example, species that independently evolved social parasitism, resulting in the loss of important social traits, showed a convergent loss of ORs, particularly within the 9-exon subfamily (63, 64). As such, it was anticipated that glomeruli number (especially in the T_B_ cluster) would also show correlated evolution with social traits. Hints of this association within the AL have also been reported, with closely related species of Dolichoderus ants exhibiting a higher number of T_B_ glomeruli with larger colony size (46). However, at a broader taxonomic scale, variation in the T_B_ cluster is not significantly associated with any of the social traits we measured, including colony size. Moreover, our ancestral trait reconstruction predicts a high number of T_B_ glomeruli in the MRCA of ants, contradicting previous suggestions of a specific expansion of the CHC-related subsystem in ants (11, 37). These results rather align with later studies showing a very high number of 9-exon ORs in non-social apoid wasps outside the Formicidae family (65), suggesting that a well-developed CHC-sensitive subsystem was conserved across these lineages and existed before the diversification of ants.

### Ecological correlates of olfactory adaptation in ants

When assessing the scaling relationship of AL subsystems as an indicator of relative investment in perceiving distinct sets of olfactory stimuli, we observed a general covariation suggesting that ants generally maintain an optimal balance between the T_B_ cluster and the Main-AL. However, *P. boopis* and Dorylinae species exhibit a notable deviation from this trend, characterized by a disproportionately high number of T_B_ glomeruli. This drastic expansion has been linked with predatory myrmecophagy, nomadism and cyclical reproduction within the Dorylinae (12). In contrast, *Pseudomyrmex* are herbivorous and omnivorous ants, residing in moderately sized colonies (66), predominantly in arboreal environment, which does not align with these interpretations.

Testing ecological factors in phylogeny-controlled GLMMs, we found evidence of a potential role of CHC complexity in influencing the relationship between the T_B_ cluster and the Main-AL. Species with a lower diversity of CHC classes in their chemical profiles exhibit a higher number of glomeruli in the T_B_ cluster. This correlation implies that the sophistication of the olfactory system may compensate for a reduced discriminative value in chemical signatures, thereby enabling accurate recognition among a smaller set of informative compounds within the CHC profile. Our behavioral experiments further support this hypothesis by demonstrating that despite variation in the number of glomeruli in the T_B_ cluster, different species perform similarly in simple nestmate discrimination tasks. Therefore, the complexity of CHC profiles likely interacts with the neuronal investment in the T_B_ cluster to maintain consistent behavioral responses in nestmate recognition, despite interspecific variation in colony-specific CHC profiles. However, it is important to note that CHC complexity exhibited a relatively low explanatory power in our analyses. Hence, in addition to other ecological factors, future research should therefore explore how the complexity of sensory systems and the chemical environment (CHC profile, but also nest and food odors) interact to drive adaptations within communication systems and influence social evolution.

Thus, ants’ remarkable evolutionary radiation, which led to a wide array of social structures and life history traits, was supported by an already complex olfactory system. In most cases, this evolution maintained a strict balance between the main-AL, responsible for detecting general odorants, and the T_B_ cluster, specialized in recognizing social cues.

## Materials and Methods

Fourteen ant species from eight subfamilies were studied, with nine species obtained from lab rearing and five collected in the wild. In all cases, ant workers were anesthetized on ice before heads and antennae were separated. Bodies were stored in 95% ethanol. To investigate phenotypic variation of the olfactory system, antennae were preserved in 2.5% glutaraldehyde at 4°C, and brains were dissected and preserved in methanol, following procedures in SI methods. In all but one species, antennal sensilla were studied using scanning electron microscopy, with standard sample preparation and scanning procedures provided in SI methods. Antennal lobe neuroanatomy was investigated through immunostaining and confocal laser scanning microscopy of stained tissues, as described in SI methods. Behavioral tests on *L. niger, M. barbarus*, and *F. fusca* were conducted to assess nestmate recognition performances, with behaviors recorded and scored for aggression, following protocols in SI methods. Chemical analyses of CHC profiles were performed via gas-chromatography coupled with mass-spectrometry (GC-MS), as described in the SI methods. Data analysis was performed using R and established statistical packages, with phylogenetic comparisons and evolutionary analyses exploring relationships between antennal morphology, neuroanatomy, chemical complexity, and behavior. All methods and statistical tests are detailed in SI methods.

## Acknowledgments

We gratefully acknowledge Fabrice Savarit for providing access to *Ectatomma tuberculatum*, and Jonathan Romiguier for supplying the time-calibrated phylogeny of ants. We also thank Chloé Leroy for assistance with the chemical analysis (GC-MS). We are grateful to the Wolfson Bioimaging Centre, University of Bristol, and Judith Mantell for their expertise in SEM imaging. We thank the Smithsonian Tropical Research Institute of Barro Colorado (Panama) for granting access to their facilities and permitting the collection of wild species. This research was supported by funding from the Leverhulme Trust (RPG-2019-287) and the French National Research Agency (ANR-20-CE02-0012).

## Competing Interest Statement

The authors declare no competing interest.

## Author Contributions

Conceptualization: A.C., P.d.E., S.H.M., J.C.S. Methodology: A.C., E.H.D., P.d.E., S.H.M., J.C.S. Investigation: S.M., A.C., E.H.D., N.B. Analysis: S.M., A.C., E.H.D. Visualization: S.M., A.C., E.H.D. Funding acquisition: A.C., P.d.E., S.H.M., J.C.S. Project administration: P.d.E., S.H.M., J.C.S. writing – original draft: S.M., A.C. Writing – review and editing: S.M., A.C., P.d.E., S.H.M., J.C.S.

## Notes

### Competing Interest Statement

The authors have declared no competing interest.

