## Supplementary information for "Ancestral complexity and constrained diversification of the ant olfactory system"

* contributed equally

#### This PDF file includes:

**1. Supplementary methods**

1.1 Species collection and sampling

1.2 Scanning electron microscopy

1.3 Immunohistochemistry

1.4 Confocal imaging

1.5 Image processing and 3D reconstructions

1.6 Behavioral experiments

1.7 Chemical analysis of CHC profiles

1.8 Statistical analysis

**2. Supplementary figures and tables**

2.1 Distribution of basiconic sensilla across ant species

2.2 Anatomy of the antennal lobe

2.3 Evolutionary dynamics of the ant antennal lobe

2.4 Influence of ecological and social factors on AL adaptation

2.5 Quantification of olfactory traits

2.6 Social and chemical predictors of antennal lobe investment

2.7 Legend for Supplementary dataset

**3. Supplementary results**

3.1 Comparative morphology of basiconic sensilla

3.2 Relationship of basiconic sensilla with AL morphology

**4. Supplementary references**

### 1. Supplementary methods

#### 1.1 Species collection and sampling

Fourteen species belonging to eight subfamilies across the phylogeny of Formicidae were chosen. Nine species were obtained from populations reared in captivity at the Laboratory of Experimental and Comparative Ethology (University Sorbonne Paris Nord, France): *Neoponera apicalis*, *Lasius niger*, *Cataglyphis cursor*, *Formica fusca*, *Camponotus aethiops*, *Ectatomma tuberculatum*, *Messor barbarus*, *Aphaenogaster senilis*, *Atta sexdens*. Five species were collected in the wild (under permit n° PA-01ARB-083-2022), on Barro Colorado Island (Republic of Panama): *Paraponera clavata*, *Eciton burchellii*, *Azteca trigona*, *Pseudomyrmex boopis*, *Cephalotes atratus*. All collected individuals were workers. Two wasp species serving as outgroups were included in our study: *Ampulex compressa* females were obtained from Aquazoo Löbbecke Museum (Düsseldorf, Germany) and *Polistes dominula* females were caught in Gif-sur-Yvette (France).

Individuals were anesthetized by cold exposure on ice before the separation of heads and antennae. The ant bodies were preserved in 95% ethanol, while the antennae were stored in a fixative solution containing 2.5% glutaraldehyde for a minimum of 7 days at 4°C. The brains were dissected out in HBS solution (150 mM NaCl, 5 mM KCl, 5 mM CaCl2, 25 mM sucrose, 10 mM HEPES, pH 7.4) and subsequently fixed in zinc-formaldehyde (4% formaldehyde; 18.4mM ZnCl2; 135mM NaCl; 35mM sucrose) for 16 to 20 h at room temperature. After fixation, brains were washed three times in HBS (10 min each), followed by dehydration in Dent's mixture (80% methanol; 20% dimethylsulfoxide) for 2 hours. Subsequently, they were transferred to a storage solution of methanol and maintained at -20°C for several months until further processing.

Behavioral experiments were carried out on three species: *Lasius niger*, *Messor barbarus* and *Formica fusca*. We used three queen-right colonies per species. *L. niger* colonies were collected in 2021 in Île-de-France, France, *M. barbarus* were collected in 2018 in Argelès-sur-Mer, France and *F. fusca* were collected in 2022 from the forest of Ermenonville, France. All colonies were kept in the laboratory under controlled conditions (25°C; natural light-dark cycle, 40% humidity). Colonies were housed in plastic boxes (25 cm x 18 cm x 9.5 cm) coated with Fluon ©. They were provided with a mixture of honey, apples and crickets twice a week, while water was available *ad libitum*.

#### 1.2 Scanning electron microscopy

To visualize the structure of ants' antennae, the fixative solution (2.5% glutaraldehyde) was removed from the antennae through three consecutive 0.1M PBS washes (for 10 min each). Subsequently, the samples were dehydrated using increasing ethanol concentrations (50%, 70%, 90%, 95%, and 3 x 100%, for 10 min each) and air dried at room temperature under a desiccator. The antennae were mounted on aluminum stubs with double-sided sticky tape, ensuring a 90° angle formed by the scape and pedicel to enable scanning of both a ventral and a dorsal antenna side from a right and a left antenna. The samples were gold-coated with a 10 nm thickness using a Quorum Q150RES sputter coater. Observations were conducted using a Zeiss Evo 15 scanning electron microscope in a high vacuum environment, utilizing secondary electron and backscatter detectors. Each flagellomere was scanned at 15 kV and 80 pA with a 7 mm working distance. High-magnification images of individual sensilla were captured at 10 kV and 100 pA.

##### 1.3 Immunohistochemistry

Unless otherwise specified, the following procedure was carried out at room temperature. Stored brains (see 1.1 Species collection and sampling) were brought back to room temperature and rehydrated progressively in a series of decreasing methanol concentrations (90%, 70%, 50%, 30% and 0% in 0.1M Tris buffer, pH 7.4, 10 min each). Brains were then incubated with collagenase-dispase (1 mg.mL$^{-1}$) in 0.01M PBS for 1 h. After two washes in PBS (10 min each), samples were permeabilized in PBS containing 2% Triton X-100 for 10 min. Then, they were blocked for 2 h in a solution of 5% normal goat serum (NGS: G9023, Sigma-Aldrich) and 0.2% Triton X-100 in PBS, hereafter referred to as PBSt-NGS. A rabbit anti-serotonin antibody (rabbit anti-5-HT; 20080; Immunostar; RRID: AB_572263), at a 1:2000 dilution in PBSt-NGS was applied for 7 days at 4°C under constant agitation, to stain serotonergic neurons. After three washes in PBSt (30 min each), brains were incubated in Cy3-conjugated goat anti-rabbit antibody (Cy3 goat anti-rabbit IgG: 111-165-144, Jackson ImmunoResearch, RRID: AB_2338006) and Alexa Fluor 488 Hydrazide (A10436; Thermo Ficher Scientific) diluted respectively at 1:200 and 1:1000 in PBSt-NGS for 7 days at 4°C, under agitation. Brains were then dehydrated through a series of increasing glycerol concentrations (1%, 2%, 4%, for 2 h each; 8%, 15%, 30%, 50%, 60%, 70% and 80%, for 1 h each) in Tris buffer, pH 7.4, with 1% dimethylsulfoxide. This was followed by five baths with 100% ethanol (10 min each). The brains were then immersed in methyl salicylate for a minimum of 3 days at 4°C before undergoing confocal imaging.

##### 1.4 Confocal imaging

Brains were mounted in the wells of aluminum slides, sandwiched between two microscope coverslips, using fresh methyl salicylate as the mounting medium. Scanning of the antennal lobes was performed using a laser-scanning confocal microscope (Zeiss LSM 700 microscope, Jena, Germany, RRID: SCR_017377) equipped with a water immersion objective (20× plan-apochromat 1.0 NA). AlexaFluor488 and Cy3 dyes were respectively excited with 555 nm and 488 nm wavelengths using solid state lasers. Antennal lobes were scanned with a resolution of 1024 x 1024 (x,y) pixels, resulting in pixel sizes ranging from 0.240 x 0.240 µm in *L. niger* to 0.521 x 0.521 µm in *P. clavata*. The entire neuropil was scanned at 1 µm intervals along the depth (z-axis).

##### 1.5 Image processing and 3D reconstructions

Scanning electron microscope images were imported in DotDotGoose software to label and count the number of basiconic sensilla (Ersts, P.J. DotDotGoose 1.6.0; American Museum of Natural History, Center for Biodiversity and Conservation). The antennal surface was assessed by measuring the length, proximal, and distal radius of each flagellomere (Fig. S1B) using ImageJ (RRID: SCR_003070) and calculating the visible surface as half the surface of a truncated cone: $S = \pi a(R + r)\ /2,$ with $a = \sqrt{h^2 + (R - r)^2}$; R being the larger radius, r the smaller radius, h the height and a the apothem of the cone.

Confocal image stacks were imported in Amira software (AMIRA 5.4.3, VSG, Berlin, Germany; RRID: SCR_007353) to perform three-dimensional analysis. Glomeruli were counted by identifying their extremities along the z-axis and using the Interpolate function to label each glomerulus in all the images where it appears. The organization of the AL into several glomerular clusters was already shown in ants (*Camponotus floridanus* (1); *Camponotus japonicus* (2); *Atta vollenweideri* (3)). Most of these clusters assemble in a large sphere around a central hub. However, these studies also reported a T6 cluster gathering tightly packed and small-sized glomeruli in the dorso-rostral region outlining a secondary glomerular sphere that flanks the rest of the AL. This T6 cluster was later named '$T_B$ cluster' in an attempt at inter-species standardization (4-6). Accordingly, glomeruli were attributed either to the '$T_B$ cluster' or to the 'Main-AL' subregions, based on distinctions in serotonin innervation and aforementioned analogous morphological features. The volume of each subregion was determined by segmenting the volume occupied by all the glomeruli belonging to these regions every five images, followed by interpolation to create a 3D model from which the volume could be extracted.

In the evolutionary analyses of glomerular counts, the tree was rooted with three outgroup species: *Ampulex compressa*, *Polistes dominula* and *Apis mellifera*. For the first two, glomeruli were counted as for ant species. For *Apis mellifera*, in which the actual presence of a $T_B$ cluster is anatomically unclear (4, 7), the number of 9-exon OR (42, (8)) was used as a proxy. The models were also run with alternative neuroanatomical values (9), including the number of T3b glomeruli (12) and T3 glomeruli (77), yielding roughly similar predictions on ancestral states.

##### 1.6 Behavioral experiments

To investigate possible behavioral differences in nestmate recognition abilities related to antennal lobe neuroanatomy, we focused on three species: *L. niger, M. barbarus* and *F. fusca* as they showed pronounced differences in serotonergic innervation and number of glomeruli. To measure nestmate discrimination ability, we conducted a discrimination task in which a single worker was simultaneously presented with two individuals from the same species, a nestmate and a non-nestmate. Individual ants were placed in fluon coated plastic arenas (height = 2.3 cm, diameter = 5 cm). The bottom was covered by a filter paper (diameter = 55 mm; Whatman), which had been placed in the test ant's colony for 24 hours prior to the beginning of experiments. This was done so that the test ant would perceive the odors of its own nest and would be more likely to engage in nest discrimination behaviors (10). The filter paper was changed after each trial. At the beginning of the test, a single ant was placed in the center of the arena and held in a small plastic fluon-coated cylinder (diameter = 2 cm, height = 2.3 cm) for 3 min, to acclimatize to the new environment. Two stimulus ants were placed in the same arena, 2.3 cm from the holding cylinder and 2 cm apart from one another. One of these stimulus ants came from the same colony as the test ant ('nestmate'), while the other originated from a different intraspecific colony ('non-nestmate'). For all colonies of the same species, the non-nestmates always originated from the same non-nestmate colony. On the day of experiments, both stimulus ants were killed by freezing and then warmed up at ambient temperature for 5 min prior to the start of the experiment. This was done to prevent the behavior of the stimulus ants from influencing that of the test ant (10). The position (right or left) of the nestmate and non-nestmate was randomized over trials.

After the 3-min acclimatization, the test ant was released from its holding tube and its behavior towards each stimulus ant was recorded using a SONY Handycam FDR AX-33. Each trial lasted 3 min during which the duration and occurrence of antennation, mandible opening and biting were recorded (10, 11). Only the most aggressive behavior of the test ant was recorded at any given time. The different behaviors were scored so that the most aggressive behaviors were assigned the highest score (0 = antennation; 1 = mandible opening; 2 = biting). These scores were used to calculate an aggression index by multiplying each aggression score with the time (in seconds) the test ants engaged in the respective behavior, and then normalizing the sum of the three resulting values with the total interaction time (sum of all behaviors) (11), using the following formula: $AI = \frac{\sum_{i=1}^{n} a_i . t_i}{T}$, Where ai and ti are the aggression score and total duration of each action respectively and T is the total interaction time across all behaviors. All videos were analyzed using the software Ethoc 1.2 (CRCA, Université Toulouse III). We tested 10 ants per colony, totaling 30 ants per species.

##### 1.7 Chemical analysis of CHC profiles

Data on chemical profiles were obtained from the literature for the following species: *E. tuberculatum* (12), *C. aethiops* (13), *C. cursor* (14), *A. sexdens* (15), *E. burchellii* (16), *N. apicalis* (17), *A. senilis* (18) and *P. clavata* (19). For the species *F. fusca*, *L. niger* and *M. barbarus*, the chemical composition of the CHC profile was determined by chemical analysis (GC-MS). Ants were killed by freezing them in individual tubes at − 20 °C. The extracts of their CHCs were obtained by dipping each ant separately in 50 µL of pentane (≥99%, HPLC grade, Sigma-Aldrich) supplemented with an internal standard (*n*-$C_{21}$ at 5 ng/µL) for 10 min. We then injected 2 µL of the extract into an Agilent 7890A gas chromatograph (GC), equipped with an HP-5MS capillary column (30 m × 0.25 mm × 0.25 µm) and a split–splitless injector, coupled to an Agilent 5975C Inert XL mass spectrometer (MS) with 70 eV electron impact ionization. The carrier gas was helium at 1 mL/min. The temperature program was as follows: an initial hold at 70 °C for 1 min, then 70–180 °C at 30 °C/min, then 180–320 °C at 5 °C/min then hold at 320 °C for 5 min. Hydrocarbons were identified by their retention times, mass spectra (diagnostic ions) and comparison with standards and published spectra.

##### 1.8 Statistical analysis

All statistical analyses were performed in R version 4.1.2 using R studio (20). Antennal data were analyzed using two-way ANOVA for segment, species and segment/species interaction effects. Interspecific variation in AL volumes and numbers of glomeruli were assessed using non-parametric Kruskall-Wallis tests. Correlation analyses between various AL data and between AL data and sensilla number were performed with Pearson tests. Ancestral state and evolutionary rate of glomeruli number were inferred using respectively the fastAnc and multirateBM functions from the R package phytools version 1.9.16 (21). The branch lengths of the phylogenetic tree were determined for eight species, each corresponding to its respective species or genus based on a recently published phylogeny (22). The species *N. apicalis*, *E. burchellii*, *A. trigona* and *C. atratus*, which were absent from this phylogeny, were substituted with their closest monophyletic species in this dataset, as this does not alter the tree structure. Thus, *N. apicalis* was substituted by *Pachycondyla rufipes* (formerly *Pseudoneoponera* *rufipes*), *E. burchellii* by *Leptanilloides erinys*, *A. trigona* by *Linepithema humile* and *C. atratus* by *Pheidole megacephala*. Without close substitutable species in the published phylogeny (22), the branch length leading to *Cataglyphis cursor* was estimated using its divergence time from *Formica sp.* (23), while the one leading to *A. senilis* was estimated using its divergence from *M. barbarus* (24). For these analyses, we included three outgroup species *Ampulex compressa*, *Polistes dominula* and *Apis mellifera*, all being present in the recently published ant phylogeny (22). Evolutionary rate analysis was performed by generating models for both $T_B$ cluster and Main-AL and by calculating the residuals from the linear regression between the extracted rates along the branches of the $T_B$ cluster versus the Main-AL. To assess how the relationship between the number of glomeruli in the $T_B$ cluster and in the Main-AL evolved, we ran a series of phylogenetic generalized linear models (GLMMs) with gaussian distributions using the R package MCMCglmm v 2.3271. The tested factors included ants' foraging mode, polygyny, colony size and the complexity of their CHC blend. These factors were tested individually to assess for a potential effect. The complexity of CHC blends was assessed using the Shannon index formula $Si = \sum_{i=1}^{N} pi \times \log pi$, a metric classically employed to quantify the diversity of ecological systems (25). In this context, Si, the Shannon index of complexity within a CHC blend, is dependent on N, the total number of different classes, and pi, the proportion of compounds belonging to a specific class (i) relative to the total number of compounds in the system. CHC were thus classified into 5 classes according to their chemical structure, i.e., alkane, alkene, methyl-alkane, dimethyl-alkane and trimethyl-alkane.

For behavioral experiments, the difference in aggression index towards nestmate and non-nestmates was analyzed separately for each species using non-parametric Kruskal-Wallis tests, which were corrected for multiple comparisons using the Benjamini-Hochberg method. Aggression index delta scores were calculated by subtracting the aggression index towards nestmates from that of non-nestmates, with positive values indicating test ants that were more aggressive towards non-nestmates and negative values indicating more aggression towards nestmates. To assess whether delta scores differed between species, data were first transformed using a Yeo-Johnson transformation to obtain a normal distribution. We then ran a linear model with species as a predictor variable and compared it to the null model using a likelihood ratio test.

#### 2. Supplementary figures

##### 2.1 Distribution of basiconic sensilla across ant species

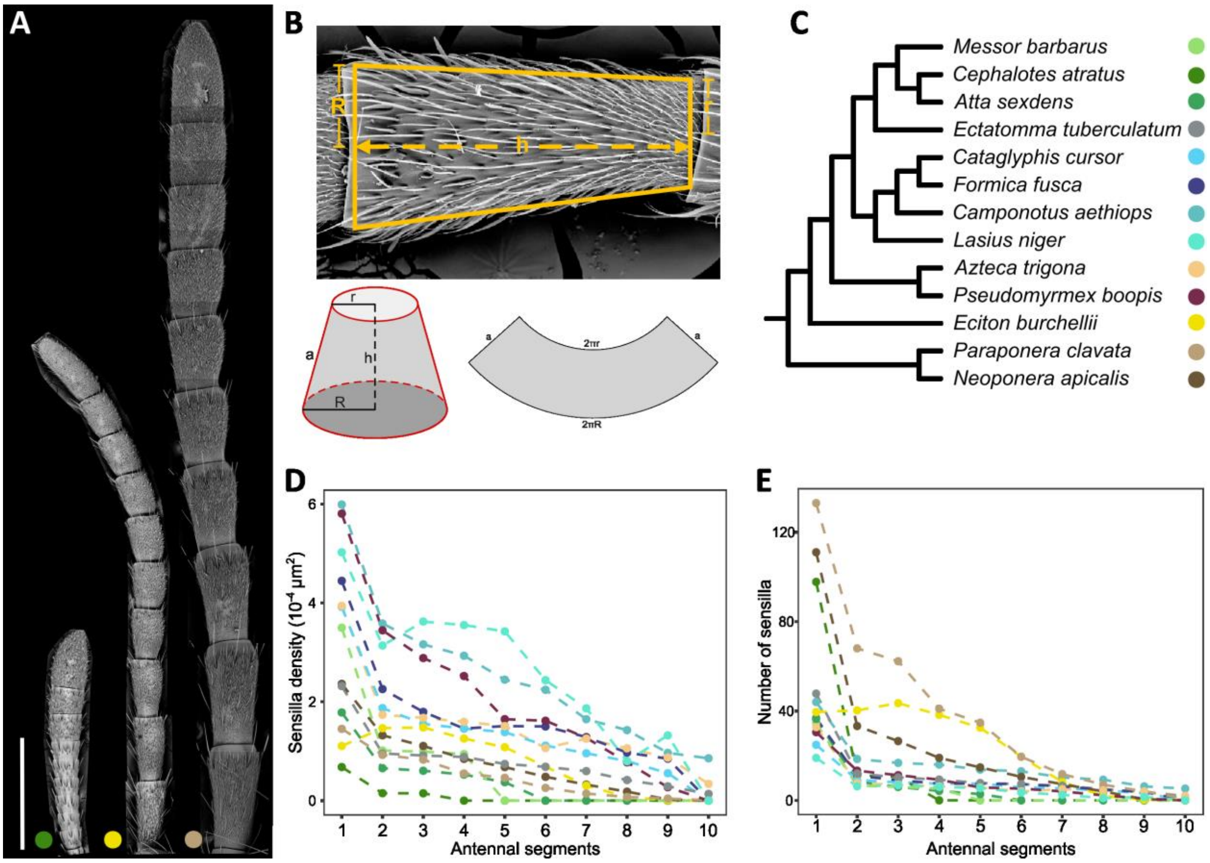


**Figure S1. Distribution of basiconic sensilla across ant species**

(A) Scanning electron micrograph of the antennae from *P. clavata*, *E. burchellii* and *C. atratus* at the same scale (scale bar = 1 mm), illustrating the variation in antenna size across ant species. (B) Antennal surface area was calculated by measuring the length (h), proximal radius (r), and distal radius (R) of each flagellomere. The visible surface was modelled as half the lateral surface area of a truncated cone (see supplementary methods for calculation). (C) Phylogeny of the sampled species, with the color coding corresponding to panels D and E (from (22-24)). (D) Density of basiconic sensilla across antennal segments in the species shown in panel C. (E) Number of basiconic sensilla across antennal segments in the species shown in panel C. The density and number of basiconic sensilla decrease progressively from the distal to the proximal regions of the antenna.

#### 2.2 Anatomy of the ant antennal lobe

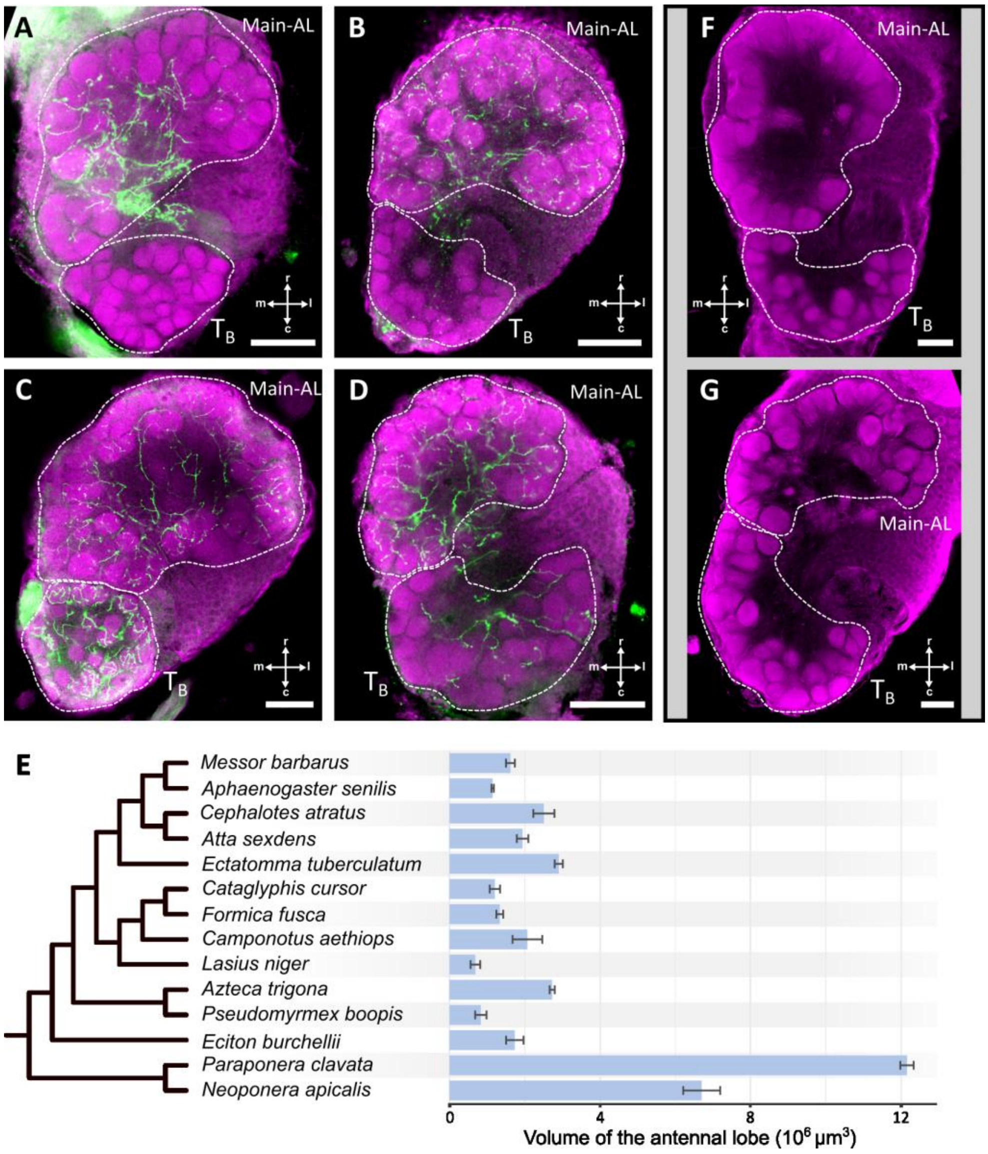


**Figure S2. Anatomy of the antennal lobe**

(A-D) Confocal optical sections of the antennal lobe in ant species that lack serotonin innervation in the $T_B$ cluster (e.g. *A. sexdens*, [A] or *C. aethiops*, [B]) and in species that show serotonin immunoreactivity in this region (e.g. *N. apicalis*, [C] or *P. boopis*, [D]). The glomeruli are stained with hydrazide conjugated dye displayed in magenta, and immunolabeled serotonergic projections are displayed in green. The AL subregions are outlined with dashed lines. (E) Bar chart showing the total antennal lobe volume across different ant species. The AL volume varies significantly between species ($\chi^2 = 38.9$, df = 14, $p < 0.001$). (F-G) For comparison, representative slices from confocal images stacks of the AL in two outgroup species, the paper wasp *Polistes dominula* (F), and the emerald cockroach wasp *Ampulex compressa* (G). All scale bars represent 50 µm (r, rostral; c, caudal; m, medial; l, lateral).

#### 2.3 Evolutionary dynamics of the ant antennal lobe

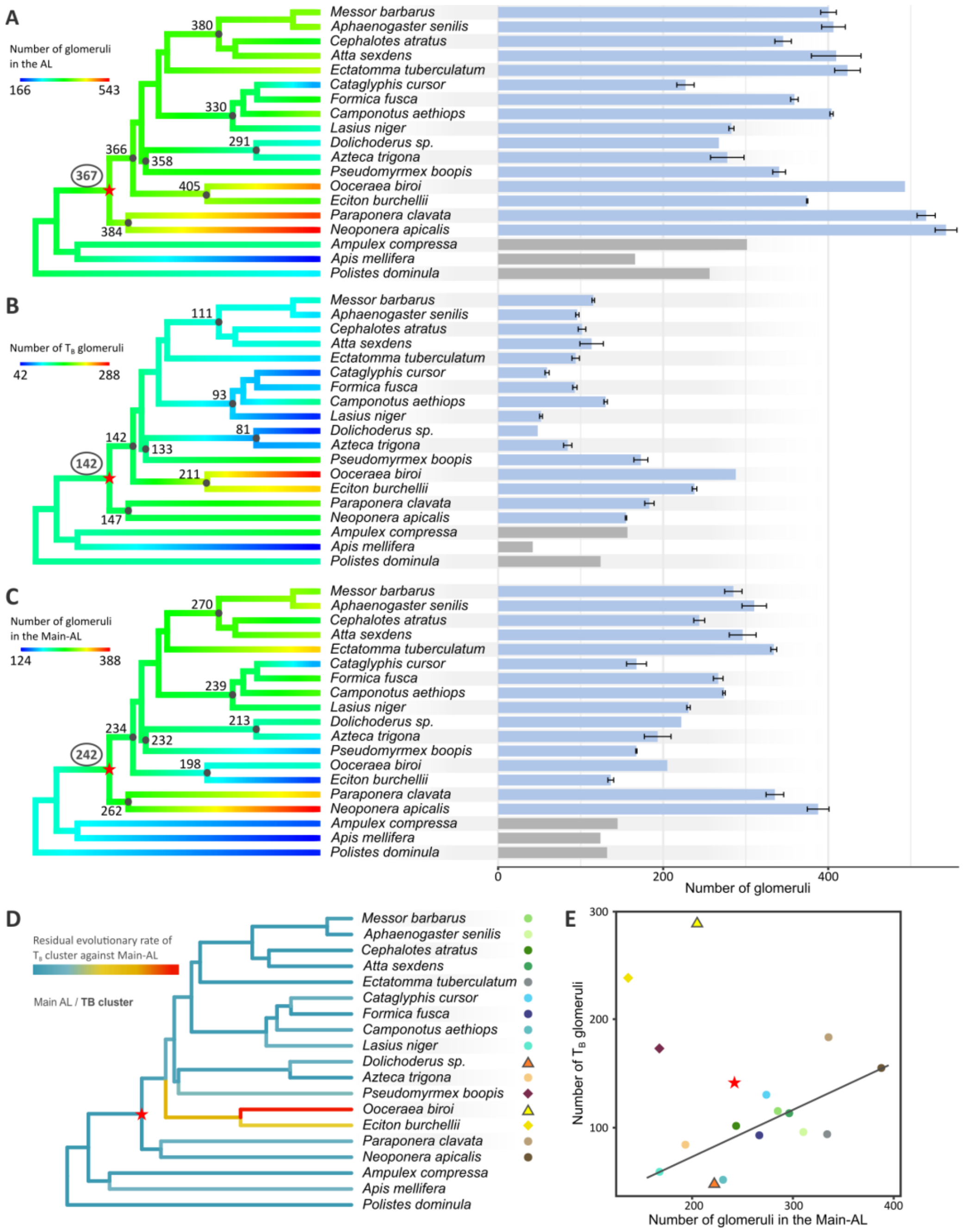


**Figure S3. Evolutionary dynamics of the ant antennal lobe**

(A-C) Left: Ancestral trait reconstruction of glomerular numbers, with values indicated at major phylogenetic nodes. Branch lengths are color-coded to show variation in glomeruli numbers. Right: bar charts display the mean (± SD) of glomeruli numbers across species. The total number of glomeruli (A), the number of $T_B$ glomeruli (B), and the number of glomeruli in the Main-AL (C) vary across species. (D) Residual evolutionary rates of glomeruli numbers in the $T_B$ cluster regressed against the Main-AL, with the branches of the phylogenetic tree color-coded to reflect these rates, including data from our sampled species as well as *Ooceraea biroi* (26) and *Dolichoderus sp.* (27) from published studies. (E) Number of $T_B$ glomeruli plotted against that of the Main-AL across different ant species. Species means are displayed with large dots, squares represent the outlier species excluded from the regression analysis and triangles the species from the literature. The red star represents the MRCA of ants. Species colors are captioned in (D).

2.4 Influence of ecological and social factors on AL adaptation

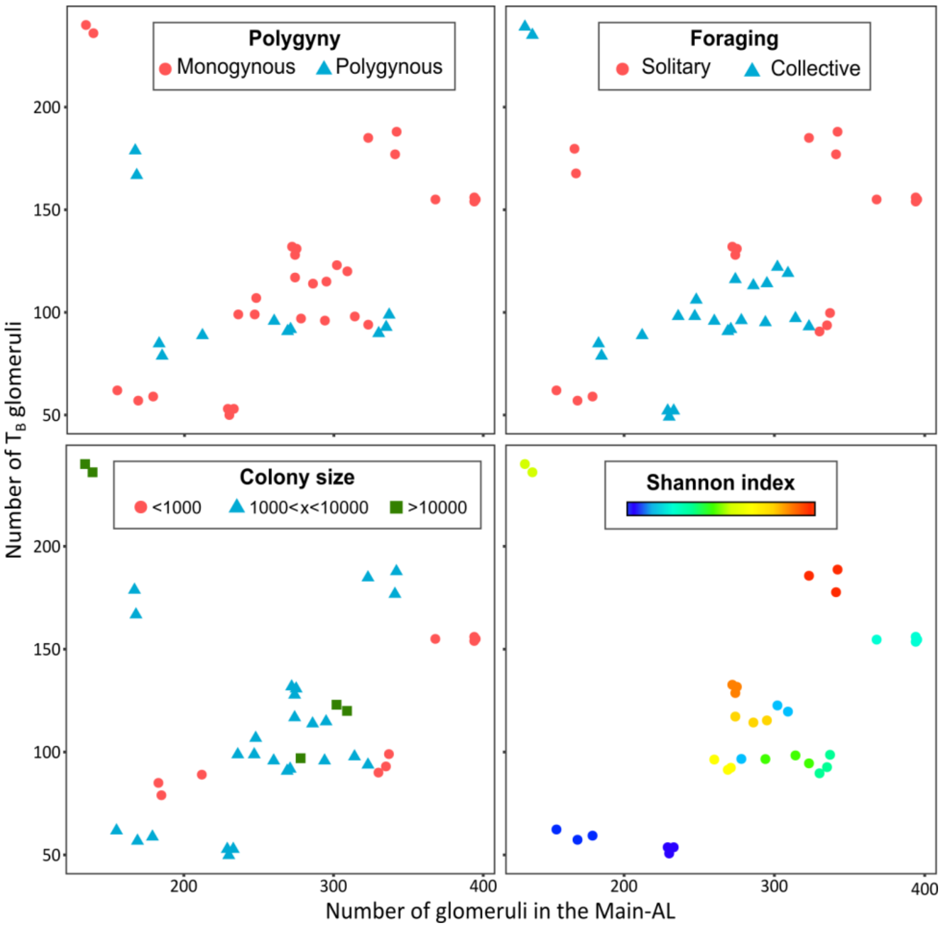


**Figure S4. Influence of ecological and social factors on investment in AL subregions**

All panels plot the number of glomeruli in the $T_B$ cluster against that in the Main-AL. Data points are color-coded based on the species' reproductive structure (monogynous or polygynous), foraging strategy (solitary or collective), colony size (3 categories), or the Shannon index measuring CHC profile complexity. Among these factors, only the Shannon index significantly predicted glomerular investment between the two subregions (*pMCMC* < 0.05).

#### 2.5 Quantification of olfactory traits

| | Sensilla | | | Whole AL | | | $T_B$ cluster | | Main-AL | |
|---|---|---|---|---|---|---|---|---|---|---|
| **Species** | **n** | **Nb. BaS.** | **Density** | **n** | **Nb. Glom.** | **Volume** | **Nb. Glom.** | **Volume** | **Nb. Glom.** | **Volume** |
| *Messor barbarus* | 5 | 52.2 ± 5.6 | 1.04E-04 ± 1.6E-05 | 3 | 400.3 ± 9.5 | 1622 ± 118 | 115.3 ± 1.5 (28.8%) | 303 ± 17 (18.7%) | 285 ± 10.5 (71.2%) | 1319 ± 101 (81.3%) |
| *Aphaenogaster senilis* | NA | NA | NA | 3 | 406.3 ± 14.4 | 1150 ± 32 | 96 ± 2 (23.6%) | 152 ± 13 (13.2%) | 310.3 ± 14.8 (76.4%) | 998 ± 19 (86.8%) |
| *Cephalotes atratus* | 4 | 119.5 ± 13.5 | 2.46E-05 ± 2.2E-06 | 3 | 345.3 ± 10 | 2512 ± 6409 | 101.7 ± 4.6 (29.4%) | 644 ± 136 (25.6%) | 243.7 ± 6.7 (70.6%) | 1868 ± 172 (74.4%) |
| *Atta sexdens* | 5 | 57.8 ± 8.8 | 6.95E-05 ± 7.6E-06 | 3 | 409.7 ± 30.1 | 1945 ± 153 | 113.3 ± 14.2 (27.7%) | 389 ± 12 (20%) | 296.3 ± 16.3 (72.3%) | 1557 ± 162 (80%) |
| *Ectatomma tuberculatum* | 4 | 111.2 ± 6.1 | 8.95E-05 ± 4.2E-06 | 3 | 428 ± 8 | 2907 ± 108 | 94 ± 4.6 (22%) | 500 ± 14 (17.2%) | 334 ± 3.6 (78%) | 2407 ± 122 (82.8%) |
| *Cataglyphis cursor* | 6 | 75.8 ± 5.8 | 1.39E-04 ± 1.2E-05 | 3 | 227 ± 10.5 | 1209 ± 137 | 59.3 ± 2.5 (26.1%) | 183 ± 7 (15.1%) | 167.7 ± 12.1 (73.9%) | 1025 ± 131 (84.9%) |
| *Formica fusca* | 4 | 93.7 ± 11.4 | 1.85E-04 ± 9.5E-06 | 3 | 359.7 ± 3.5 | 1338 ± 92 | 93 ± 2.6 (25.9%) | 253 ± 10 (18.9%) | 266.7 ± 5.9 (74.1%) | 1086 ± 84 (81.1%) |
| *Camponotus aethiops* | 4 | 153 ± 45.3 | 2.54E-04 ± 6.7E-05 | 3 | 404 ± 2 | 2074 ± 396 | 130.3 ± 2.1 (32.3%) | 443 ± 57 (21.3%) | 273.7 ± 1.5 (67.7%) | 1631 ± 343 (78.7%) |
| *Lasius niger* | 4 | 52 ± 7.0 | 3.08E-04 ± 5.1E-05 | 3 | 282.7 ± 3.1 | 690 ± 128 | 52 ± 1.7 (18.4%) | 79 ± 12 (11.4%) | 230.7 ± 2.1 (81.6%) | 612 ± 117 (88.6%) |
| *Azteca trigona* | 4 | 79 ± 4.8 | 1.68E-04 ± 7.1E-06 | 3 | 277.7 ± 20.3 | 835 ± 1125 | 84.3 ± 5 (30.4%) | 153 ± 29 (18.3%) | 193.3 ± 16.2 (69.6%) | 682 ± 123 (81.7%) |
| *Pseudomyrmex boopis* | 4 | 82.2 ± 8.4 | 2.33E-04 ± 2.6E-05 | 2 | 340.5 ± 7.8 | 1738 ± 1882 | 173 ± 8.5 (50.8%) | 706 ± 81 (40.6%) | 167.5 ± 0.7 (49.2%) | 1033 ± 152 (59.4%) |
| *Eciton burchellii* | 4 | 224 ± 64.9 | 7.60E-05 ± 2.5E-05 | 2 | 374.5 ± 0.7 | 2732 ± 2148 | 238 ± 2.8 (63.6%) | 1383 ± 60 (50.6%) | 136.5 ± 3.5 (36.4%) | 1349 ± 8 (49.4%) |
| *Paraponera clavata* | 4 | 382 ± 80.5 | 4.62E-05 ± 9.0E-06 | 3 | 518.7 ± 11 | 12168 ± 177 | 183.3 ± 5.7 (35.3%) | 3141 ± 599 (25.8%) | 335.3 ± 10.7 (64.7%) | 9027 ± 1557 (74.2%) |
| *Neoponera apicalis* | 4 | 226.7 ± 30.7 | 9.27E-05 ± 1.5E-05 | 4 | 542.8 ± 13.2 | 6777 ± 423 | 155 ± 0.8 (28.6%) | 1329 ± 207 (19.6%) | 387.8 ± 13.2 (71.4%) | 5448 ± 227 (80.4%) |

**Table S1. Mean values of quantifiable anatomical olfactory traits for each species**

For each species, the table provides the mean (± SD) number of basiconic sensilla (BaS) and their overall density, as well as the number of glomeruli and antennal lobe (AL) volumes. AL traits are detailed separately for the two subregions: the $T_B$ cluster and the Main-AL, with the proportion of each subregion relative to the total AL indicated in parentheses.

#### 2.6 Social and chemical predictors of AL investment

| Species | Polygyny | Colony size | Foraging | CHC Shannon Index | CHC class | Compounds |
|---|---|---|---|---|---|---|
| *Messor barbarus* | Monogynous (28) | 1000<x<10000 (29) | Cooperative (29) | 1 | 4 (alka, alke, me, dim) | 51 |
| *Aphaenogaster senilis* | Monogynous (30) | 1000<x<10000 (31) | Cooperative (32) | 1.17 | 4 (alka, me, dim, trim) | 34 |
| *Cephalotes atratus* | Monogynous (33) | 1000<x<10000 (33) | Cooperative (34) | NA | NA | NA |
| *Atta sexdens* | Monogynous (35) | >10000 (29) | Cooperative (29) | 1.32 | 4 (alka, me, dim, trim) | 14 |
| *Ectatomma tuberculatum* | Facultatively polygynous (36) | <1000 (37) | Solitary (38) | 1.2 | 4 (alka, alke, me, dim) | 66 |
| *Cataglyphis cursor* | Monogynous (39) | 1000<x<10000 (29) | Solitary (29) | 1.34 | 4 (alka, alke, me, dim) | 66 |
| *Formica fusca* | Facultatively polygynous (40) | 1000<x<10000 (38) | Cooperative (29) | 1.03 | 4 (alka, me, dim, trim) | 76 |
| *Camponotus aethiops* | Monogynous (41) | 1000<x<10000 (29) | Solitary (29) | 0.92 | 3 (alka, me, dim) | 73 |
| *Lasius niger* | Monogynous (42) | 1000<x<10000 (29) | Cooperative (29) | 1.37 | 5 (alka, alke, me, dim, trim) | 61 |
| *Azteca trigona* | Secondary monogynous (43) | <1000 (43) | Cooperative (44) | NA | NA | NA |
| *Pseudomyrmex boopis* | Facultatively polygynous (45) | <1000 (46) | Solitary (47) | NA | NA | NA |
| *Eciton burchellii* | Monogynous (48) | >10000 (29) | Cooperative (29) | 1.04 | 3 (alka, alke, me) | 29 |
| *Paraponera clavata* | Monogynous (49) | 1000<x<10000 (49) | Solitary (50) | 0.99 | 3 (alka, alke, me) | 59 |
| *Neoponera apicalis* | Monogynous (51) | <1000 (29) | Solitary (52) | 1.21 | 4 (alka, alke, me, dim) | 29 |

**Table S2. Ecological and colonial factors across species**

This table lists the ecological and colonial factors known for each species, which include reproductive structure, foraging strategy, colony size, and chemical traits related to each species' CHC profile. The term "monogynous" indicates species that strictly have a single queen per colony, while "polygyny" also includes secondary monogyny. CHC classes are indicated as follows: alka: alkanes, alke: alkenes, me: methyl-branched alkanes, dim: dimethyl-branched alkanes, trim: trimethyl-branched alkanes.

##### 2.7 Legend for Supplementary data file

Table presenting the results of MCMC GLMM models testing the associations between traits (e.g., glomeruli number and volume, sensilla number and density) and chemical as well as social traits. For each predictor, the table provides the posterior mean estimate, the 95% credible interval, and the pMCMC value, which indicates the significance of the predictor within the full model. The effective sample size (ESS) reflects the number of effectively independent samples from the posterior distribution. Additionally, the variance components for random effects are partitioned, showing the variance attributable to phylogeny and individuals when the corresponding factor is dropped from the model. The deviance information criterion (DIC) is reported for each full model (highlighted with grey rows) and for models where each factor is individually dropped (white rows).

#### 3. Supplementary Results

##### 3.1 Comparative morphology of basiconic sensilla

In our investigation of basiconic sensilla distribution, we observed subtle morphological variations across ant lineages. In basal species within the ponerine subfamily, the sensilla appeared stubbier, with a larger base and shorter shaft (Fig. 1A). In contrast, in more derived groups such as the Formicinae and Myrmicinae subfamilies, the sensilla had a slender appearance with a longer shaft and pointed tip (Fig. 1B-C).

Contrary to previous reports describing concave depressions at the terminal ends of basiconic sensilla in certain species (53), we found no such structures in any of the species we examined, even within the same genera as in the published study, such as *Camponotus* and *Formica*. This discrepancy might be attributed to differences in sample preparation. Specifically, the vacuum conditions during traditional SEM, combined with the lack of fixation and dehydration, can cause the collapse or deformation of delicate chitin structures, leading to artifacts such as the presence of concave tips (54). To address this, we conducted additional imaging using Environmental SEM (ESEM) on fresh, untreated antennae. As we gradually increased the chamber pressure, we observed the sensilla sinking into their sockets and eventually forming depressions at the tip. These artifacts only became apparent under high-pressure conditions, suggesting that earlier observations of concave terminals might be due to similar preparation-induced effects. These artifacts disappeared when samples were subjected to prolonged fixation in glutaraldehyde and dehydration through an ethanol series.

##### 3.2 Relationship between basiconic sensilla numbers and AL morphology

We examined the connection between basiconic sensilla and the $T_B$ cluster through straightforward correlations of numerical and volumetric traits. Both the size of the $T_B$ cluster and the mean glomerular volume within it are correlated with the number of basiconic sensilla (Fig. S5A, B; Pearson test, $t = 10.9$, $df = 11$, $p < 0.001$, $R^2 = 0.91$ and $t = 8.16$, $df = 11$, $p < 0.001$, $R^2 = 0.78$, respectively). This correlation was anticipated, as more sensilla likely imply more OSNs projecting into the glomeruli, enlarging their volume. But similarly, there is a significant correlation between both the volume of the Main-AL and the mean glomerular size with the number of basiconic sensilla (Fig. S5A, B; Pearson test, $t = 6.2986$, $df =$

11, $p < 0.001$, $R^2 = 0.86$ and t = 9.6943, df = 11, $p < 0.001$, $R^2 = 0.9$, respectively). This may be a consequence of the consistent proportionality in the number and volume of glomeruli between the two AL subdivisions. Nevertheless, the number of basiconic sensilla correlates with the number of $T_B$ glomeruli (Fig. S5C; Pearson test, t = 3.0713, df = 11, $p < 0.05$, $R^2 = 0.46$), but not with the number of glomeruli in the Main-AL (Pearson test, t = 1.2033, df = 11, p = 0.25). This suggests that species with more $T_B$ glomeruli may have more basiconic sensilla to accommodate increased OSN demand.

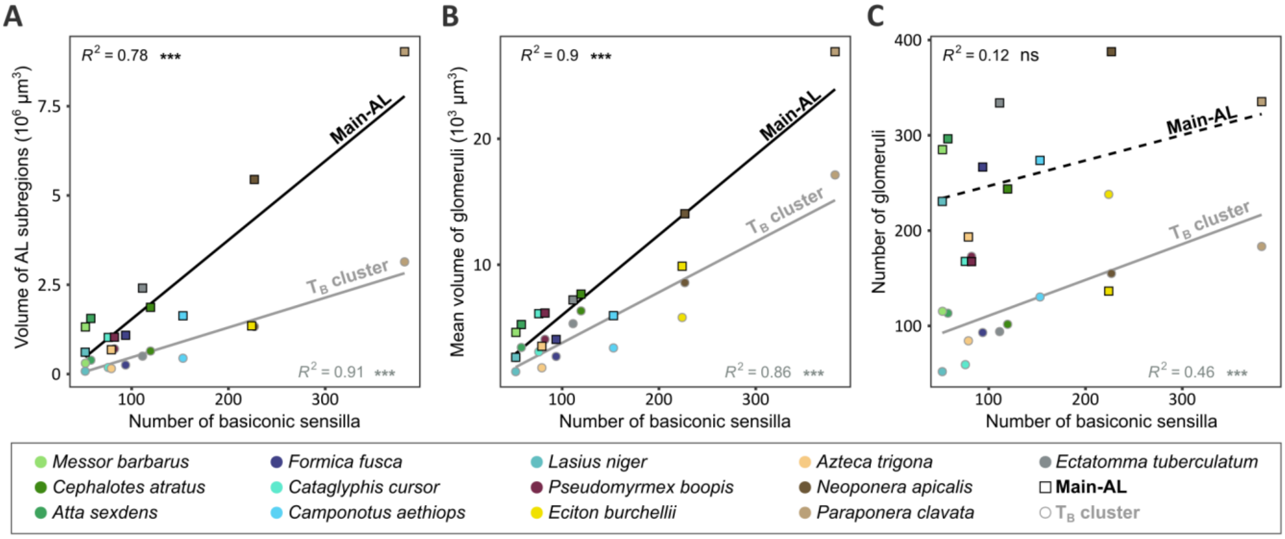


**Figure S5. Relationship between the number of basiconic sensilla and antennal lobe traits**
(A) Volume of the $T_B$ cluster (grey regression line) and the Main-AL (black regression line) plotted against the number of basiconic sensilla. (B) Mean glomerular volume in the $T_B$ cluster (grey regression line) and in the Main-AL (black regression line) plotted against the number of basiconic sensilla. (C) Number of glomeruli in the $T_B$ cluster (grey regression line) and in the Main-AL (black regression line) plotted against the number of basiconic sensilla.
